## Supplemental Information for "Commensal to pathogen switch in *Streptococcus pneumoniae* is governed by a thermosensing master regulator"

**This file contains:**

Figures S1 to S7

Table S1 to S2

**Suppl. Fig. 1**

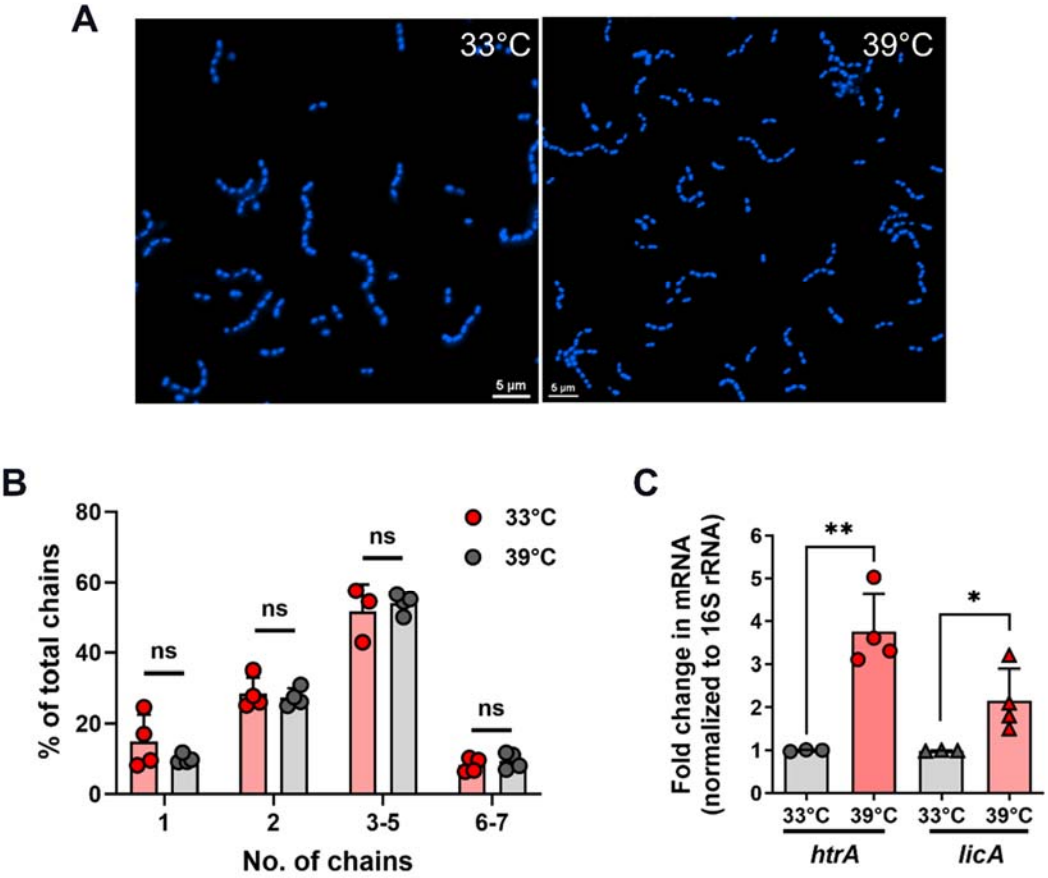


**Febrile temperature exposure doesn't affect the chain length in SPN**. **A**. Representative Immunofluorescence image displaying SPN stained with Hoest dye to quantify chain number. The temperature that they were exposed to before imaging is indicated in the left right corner. Scale bar = 5µm. **B**. Bar graph displaying quantification bacterial chain number in WT SPN exposed to 33°C and 39°C. ImageJ (Fiji) software was used to quantify chain numbers and their frequency. **C**. qRT-PCR analysis of *htrA* and *licA* transcript levels in WT SPN exposed to different temperatures. 1 µg of total RNA isolated from SPN grown at different temperatures was reverse transcribed to synthesize cDNA and equal amount of cDNA from each test sample was used for analyzing Ct values using $2^{-\Delta\Delta Ct}$ method. Transcript levels of each gene were normalized to 16S rRNA and expressed as fold change compared to 33°C. Statistical significance was assessed by two-tailed unpaired student's t-test (**B** and **C**). *P < 0.05; **P < 0.01; ***P < 0.005. Data are mean ± SEM of 3 independent biological replicates.

**Suppl. Fig. 2**

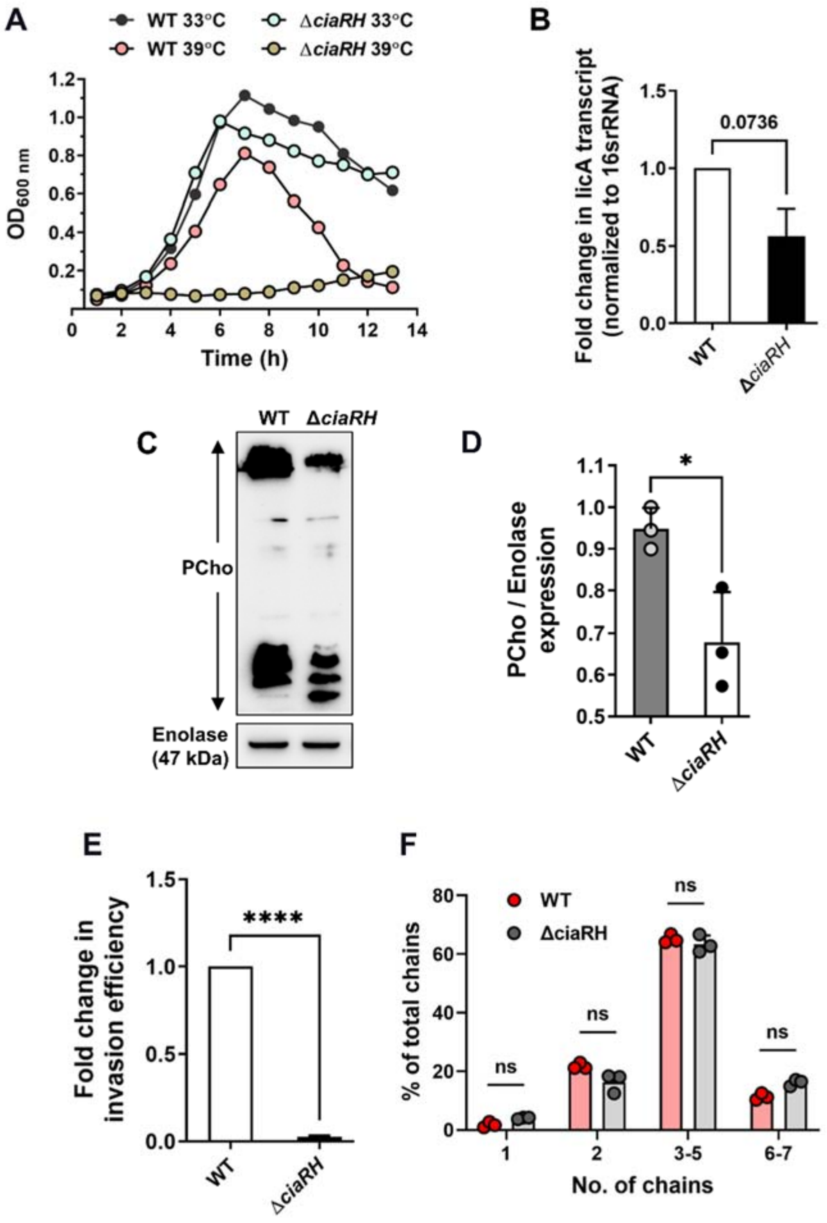


**CiaRH mutant strain of SPN demonstrates reduced PCho uptake and invasion**. **A**. Growth curve comparing growths of WT and Δ*ciaRH* SPN strain at 33°C and 39°C. **B**. Bar graph depicting abundance of *licA* transcript in WT and Δ*ciaRH* SPN strains using qRT-PCR analysis. 1 $\mu$g of total RNA was reverse transcribed to synthesize cDNA and equal amount of cDNA from each test sample was used for analyzing Ct values using $2^{-\Delta\Delta Ct}$ method. 16S rRNA transcript level was used to normalize the Ct values of each test transcripts and represented as fold change with respect to 33°C. **C**. Immunoblot demonstrating PCho levels in WT and Δ*ciaRH* SPN. Enolase was used as a loading control. **D**. Densitometric quantification of ‘C’. **E**. Fold change in invasion efficiency of WT and Δ*ciaRH* SPN in A549 cells at 37°C. Fold change was calculated as ratio of intracellular bacterial CFU of the Δ*ciaRH* to that of WT SPN. **F**. Bar graph demonstrating the chain number of WT and Δ*ciaRH* strain. ImageJ (Fiji) software was used to quantify chain numbers and their frequency. Statistical significance was assessed by two-tailed unpaired student's t-test (**D, E** and **F**). *P < 0.05; **P < 0.01; ***P < 0.005. Data are mean ± SEM of 3 independent biological replicates.

**Suppl. Fig. 3**

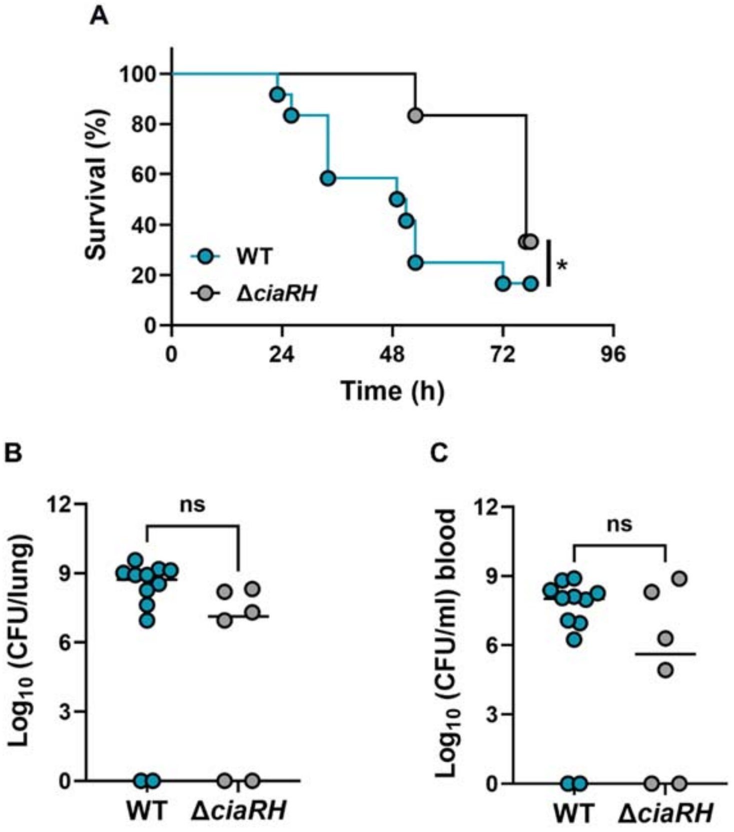


**Absence of CiaRH signaling attenuates virulence in a mouse model of invasive pneumococcal pneumonia. A.** Mice were infected intranasally with 2x106 CFU and monitored for signs of disease over 78 hours. Mice were culled when they reached pre-determined disease severity endpoints. n = 10, *P < 0.05 in Kaplan-Meier survival analysis. **B.** Bacterial burden in lungs and C. blood at time of death or, for mice that survived infection, at 78 hours post-infection. ns = not significant in Mann-Whitney test.

**Suppl. Fig. 4**

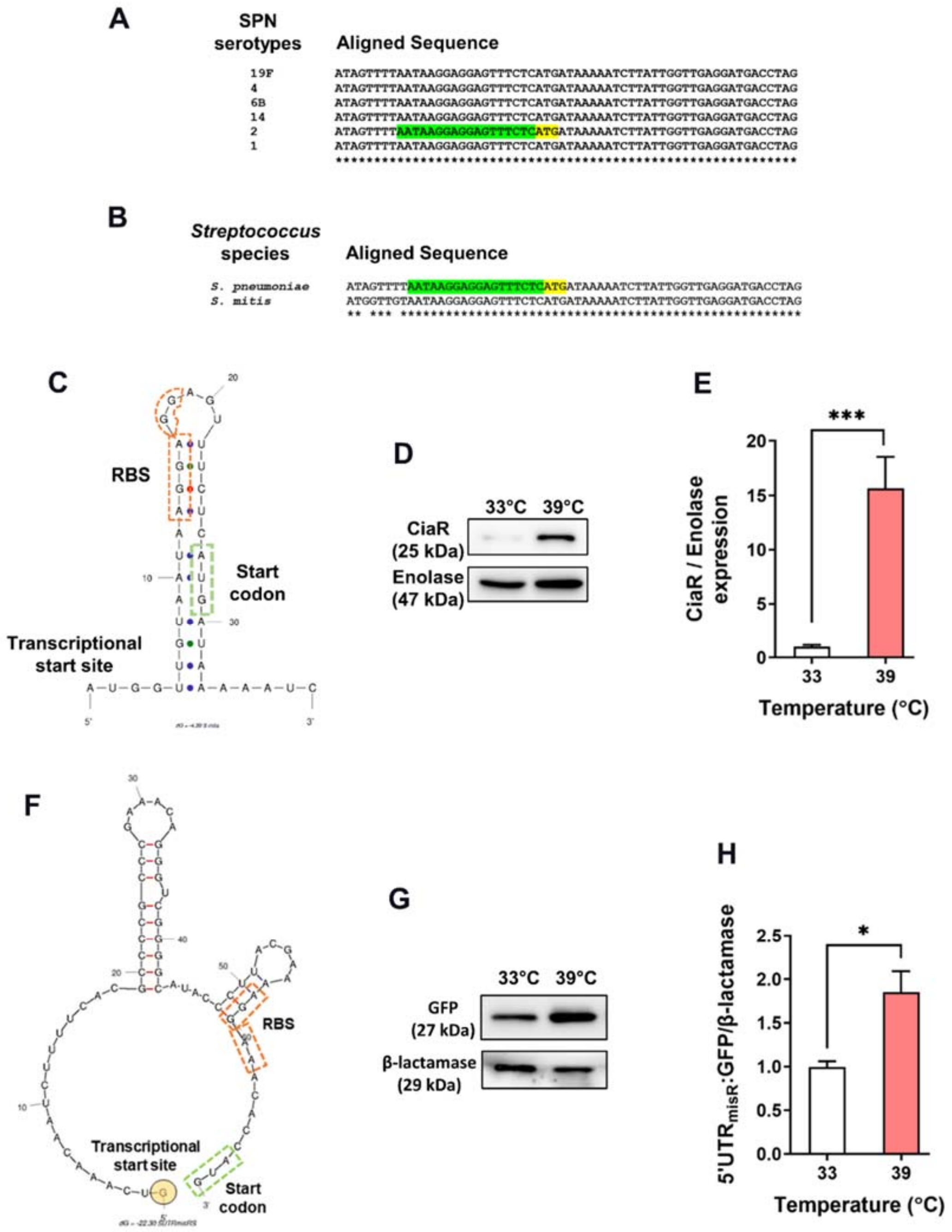


**Conserved thermosensing unit in 5'UTR of *ciaRH* shows hairpin structure**. Sequence alignment of 5'UTR *ciaRH* sequences in different SPN serotypes (**A**) and related Streptococcal species, such as *Streptococcus mitis* (**B**). Upstream region is highlighted in green while the start codon is highlighted in yellow in reference sequence. **C**. mFold® predicted secondary structure of 5'UTR of *ciaRH* of *S.mitis* marked with predicted Transcription Start Site (highlighted in yellow), Ribosome Binding Site (marked in orange) and Start codon (marked in green). **D**. Immunoblot demonstrating CiaR levels in *S. mitis* at 33°C and 39°C. Enolase served as loading control. **E**. Densitometric quantification of CiaR in 'D'. **F**. mFold® predicted secondary structure of 5'UTR of *misRS* in *Neisseria meningitidis* marked with predicted Transcription Start Site (highlighted in yellow), Ribosome Binding Site (marked in orange) and Start codon (marked in green). **G**. Immunoblot of GFP expressed under the promoter of *misRS* operon. 20 $\mu$g protein was loaded for each sample and β-lactamase served as loading control. **H**. Graph representing densitometric quantification of 'G'. Statistical significance was assessed by two-tailed unpaired student's t-test (**E** and **H**). *P < 0.05; **P < 0.01; ***P < 0.005. Data are mean ± SD of 2 independent biological replicates.

**Suppl. Fig. 5**

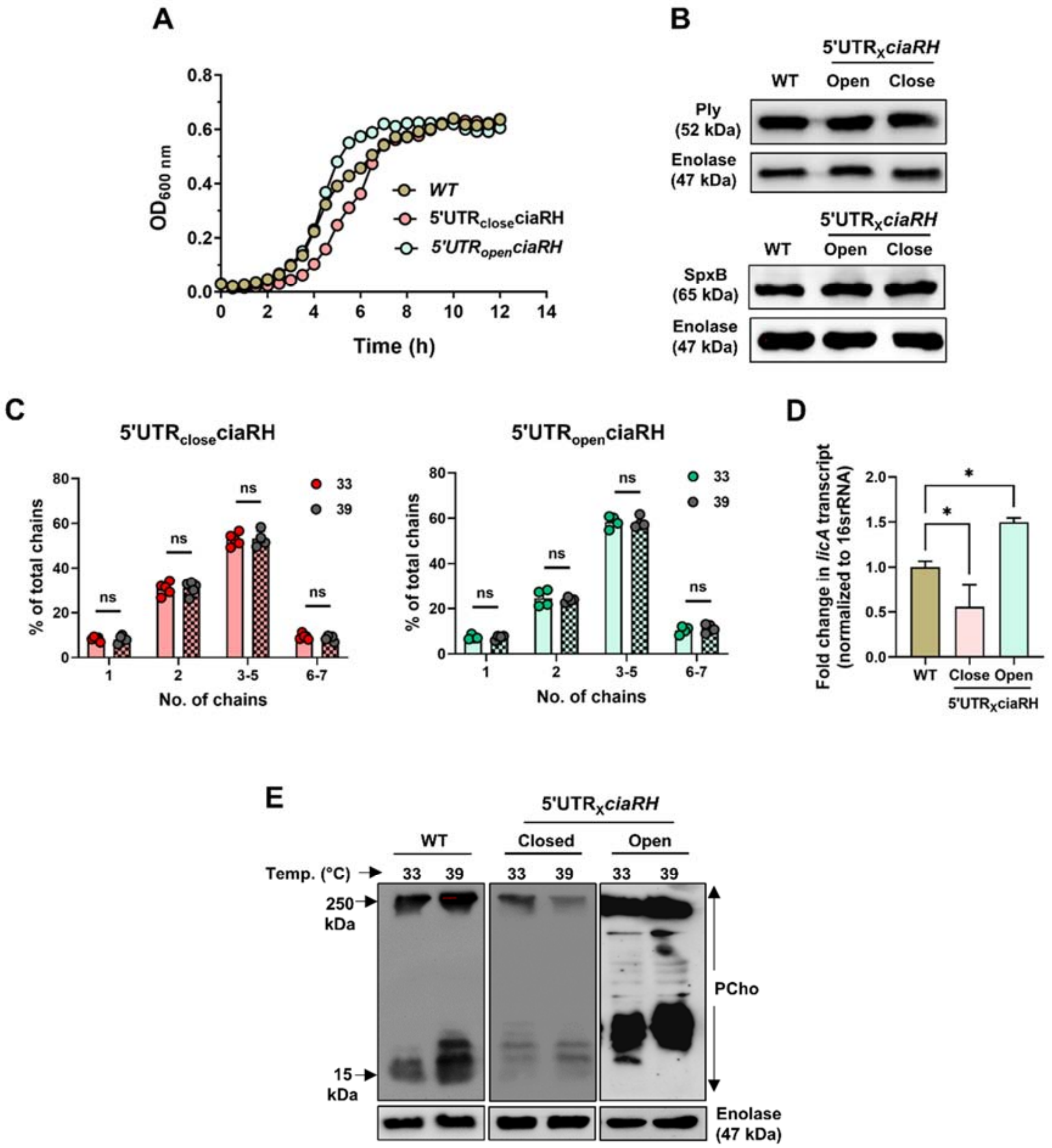


**Characterization of mutated 5'UTR *ciaRH* SPN strains**. **A**. Plot displaying growth curve of WT and 5′UTR$_{close/open}$ *ciaRH* SPN strains at 37°C. **B**. Immunoblot depicting expression of SpxB and Pneumolysin in 5′UTR$_{close/open}$ *ciaRH* strains compared to WT SPN. 20 $\mu$g of total protein was loaded in each sample and enolase was used as loading control. **C**. Bar graph quantifying chain length and their frequency in mutated 5′UTR *ciaRH* strains. Fiji® software was used for marking and quantifying the pneumococcal cells and chains. **D**. qRT-PCR analysis of *licA* transcript level in 5′UTR$_{close/open}$ *ciaRH* and WT SPN strains. 1 $\mu$g of total bacterial RNA was used for reverse transcribed to cDNA and equal amount of cDNA from each test sample was used for analyzing Ct values. 16S rRNA transcript level was used to normalize the Ct values of each test transcripts using $2^{-\Delta\Delta Ct}$ method. **E**. Representative immunoblot displaying PCho levels in WT and 5'UTR *ciaRH* mutated strains of SPN. Enolase served as loading control. Statistical significance was assessed by two-tailed unpaired student's t-test (**C**) and one-way ANOVA (Dunnett's test) (**D**). *P < 0.05; **P < 0.01; ***P < 0.005. Data are mean ± SEM of 3 independent biological replicates.

**Suppl. Fig. 6**

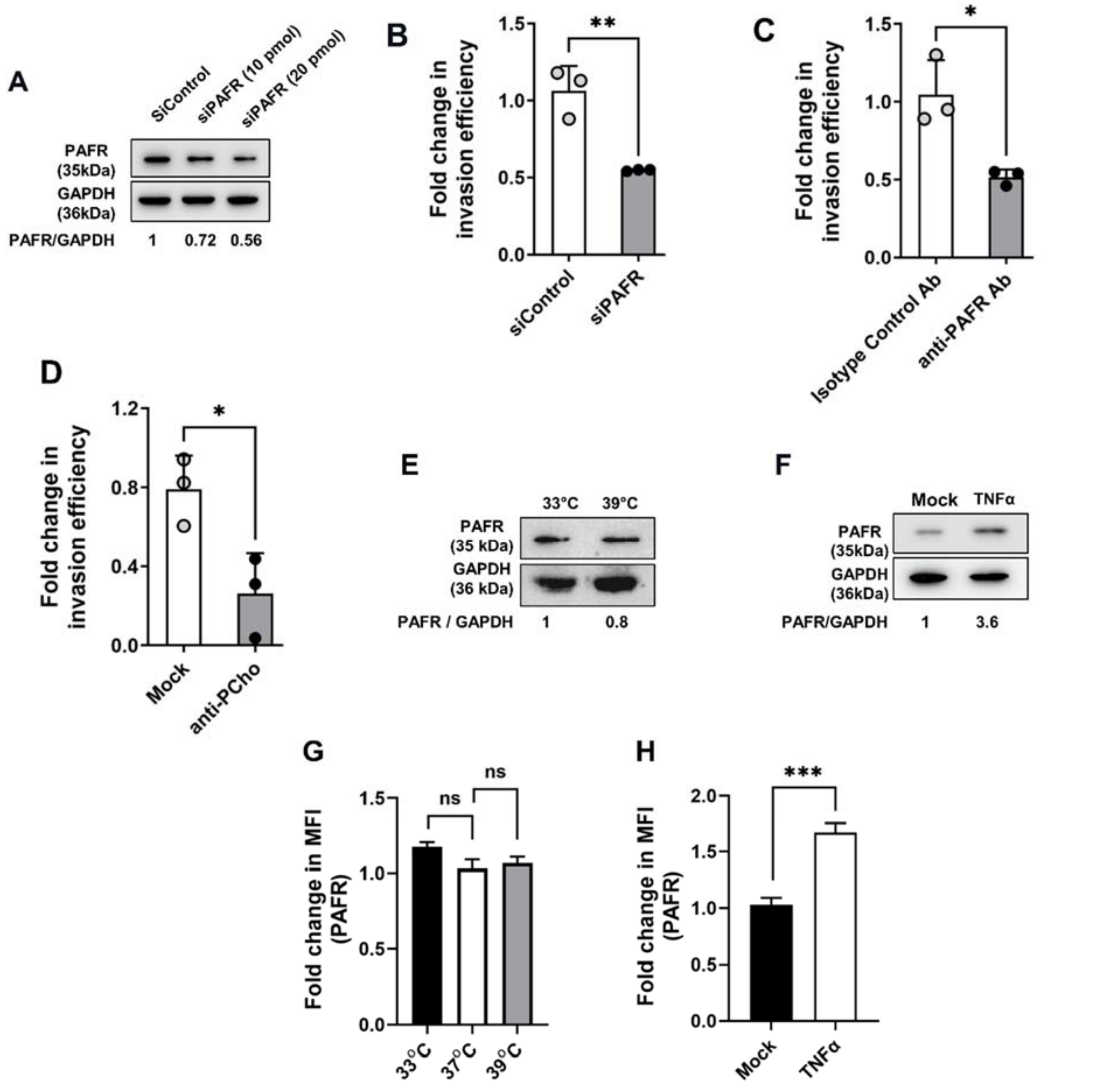


**Platelet Activating factor Receptor (PAFR) is crucial for pneumococcal uptake**. **A**. Immunoblot displaying PAFR levels in siControl and siPAFR treated A549 cells. Fold change in normalized PAFR levels with respect of GAPDH levels is mentioned below the blot. **B. C**. Graph depicting fold change in invasion efficiency of WT SPN in A549 cells treated with either siPAFR (20 pmol) for 24h **(B)**; or blocked with anti-PAFR antibody (1:1000) for 2h prior to infection **(C)**. Scrambled and isotype control antibody served as controls, respectively. Fold change was calculated as ratio of intracellular bacterial CFU of the test to that of control. **E**. **F**. Immunoblot of PAFR levels in A549 cells exposed to 33°C and 39°C **(D)** or treated with 30ng/mL TNFα **(E)**. Fold change in levels of PAFR in comparison to GAPDH in each test sample is denoted below the blot. **G**. **H**. Graph depicting fold change in MFI (Mean Fluorescence Intensity) in PAFR signal in A549 cells exposed to 33, 37 and 39°C **(D)** or treated with TNFα **(E)** as acquired by flow cytometry. Fold change was calculated with respect to A549 cells grown at 37°C (unexposed) in **D** or mock treated in **E**. Statistical significance was assessed by two-tailed unpaired student's t-test **(B, C, D** and **H)** and one-way ANOVA (Dunnett's test) **(G)**. *P < 0.05; **P < 0.01; ***P < 0.005. Data are mean ± SD of 3 independent biological replicates.

**Suppl. Fig. 7**

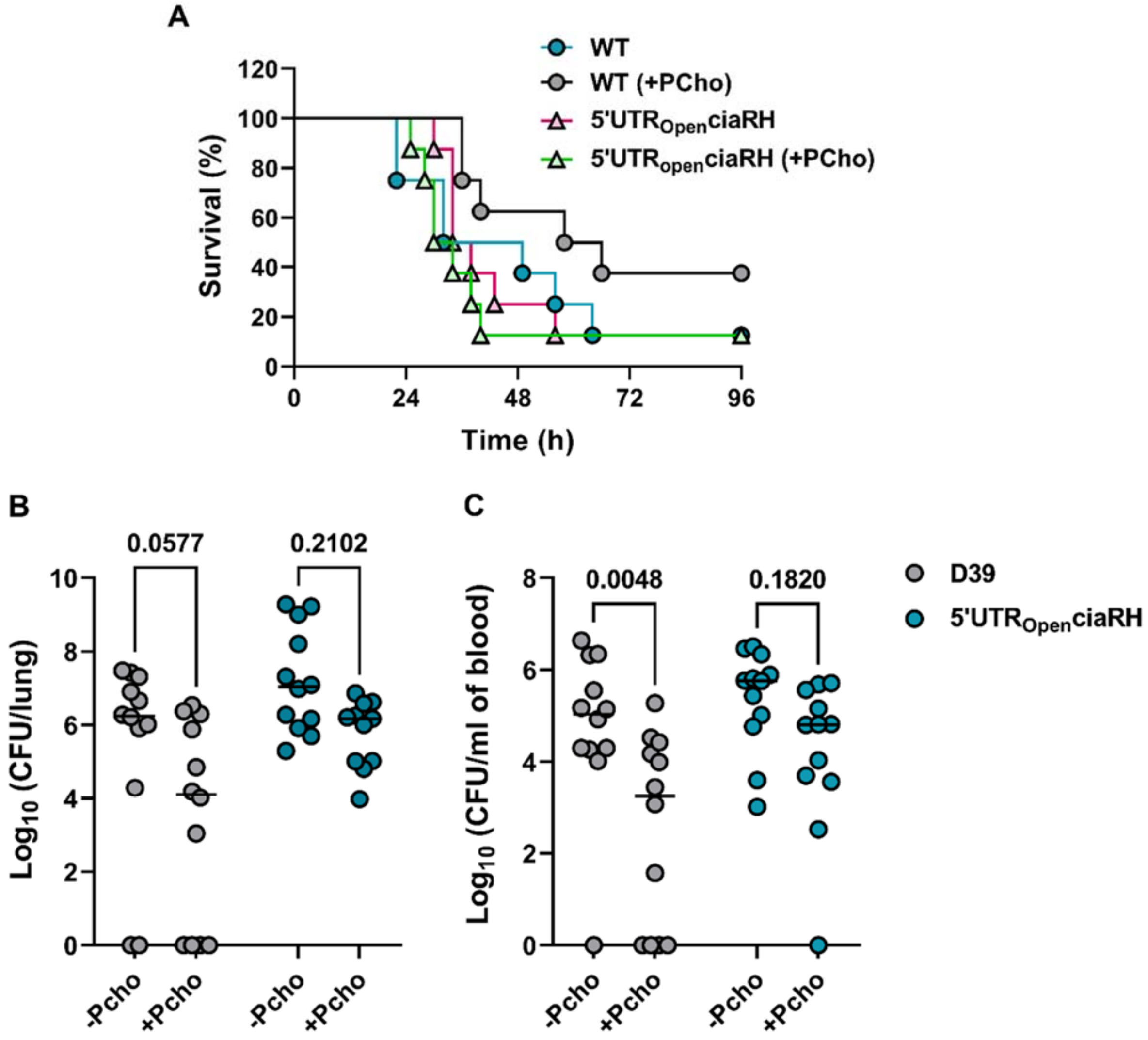


**Pre-treatment of mice with phosphorylcholine influences disease progression in pneumococcal pneumonia. A.** Mice were administered 2 mg PCho intranasally and two hours later were infected intranasally with $2x10^6$ CFU and monitored for signs of disease over 96 h. Mice were culled when they reached pre-determined disease severity endpoints. n = 10, *p < 0.05 in Kaplan-Meier survival analysis. **B-C.** Bacterial burden in lungs **(B)** and blood **(C)** at 24 h post-infection. Stattistical significance was achieved by Mann-Whitney test and mentioned in each graph.

**Table S1. List of strains**

| Strain | Source |
| --- | --- |
| *Streptococcus pneumoniae* (SPN) strain D39 (Serotype 2, encapsulated) | Prof. E. Tuomanen (St. Jude Children's Hospital, USA) |
| D39Δ*ciaRH* | This study |
| D39Δ*ciaRH*: 5'$UTR_{closed}$ *ciaRH* | This study |
| D39Δ*ciaRH*: 5'$UTR_{open}$ *ciaRH* | This study |

### Table S2. List of Primers

| Primer name | Sequence (5'-3') |
|---|---|
| GFP-F-BamHI | ATTTAAGGATCCAAAGGAGAAGAGCTGTTCACA |
| GFP-R-XhoI | GTTAGTCTCGAGTTACTTATAAAGCTCATCCAT |
| PciaR-F | GTAGGCTTCCTAATACGACTCACTATAGGGAATAAGGAGGAGT TTCTCATG |
| PciaR-R | GTTAGTCTCGAGTTACTTATAAAGCTCATCCAT |
| Peno-F | CTGACTGACGGATCCCATTTTTTACTCTCCTTATGAG |
| Peno-R | ACCGCGGTGGCGGCCGCTCTAGAGAGCTTTTTCAAGTA |
| open ciaR-F | GAATATAATGATAAAAATCTTATTGGTTGAGG |
| open ciaR-R | TCCTCCTTATTAAAACTATTATACCAAATTTG |
| closed ciaR-F | AACAAGGAGGTTCCCCCCATGATAAAAATCTTATTGGTTGAGG |
| closed ciaR-R | AAAACTATTATACCAAATTTGCCTTAAAAAAAAC |
| ciaR-F-pbsk-XhoI | TTGCATGCCTCGAGATAAGCCTAAAATAAAAAGAAAACTCAGCT ATCTCATGTAA |
| ciaR-R-pbsk-His6-EcorI | ATCGATCGAGAATTCTTAGTGGTGATGGTGATGATGCTGAACA TCTTTTAAAAGA |
| PmisR-F | GCTCTTCTCCTTTGGATCCCATGGTGTTTCCTTTTCGTAAG |
| PmisR-R | GCGGTGGCGGCCGCTCTAGACATGCAAGACATTGCAAAAA |
| 16SrRNA-F | AACCAAGTAACTTTGAAAGAAGAC |
| 16SrRNA-R | AAATTTAGAATCGTGGAATTTTT |
| Spec-R-BamHI | ATCCGGATCCAATCTGATTACCAATTAGAATG |
| specF2-BamHI | CCGCGGATCCCATATATAATCTAGAATAAAATTAAC |
| LicA-F-RT | CGATTTGGTGCCTGAAAACT |
| LicA-R-RT | ACCGGTGTTTGGTCACTCTC |
| CiaR-F-RT | GATGGAGAAGAAGGTC |
| CiaR-R-RT | GTCATAATCAGAACTGG |
| HtrA-F-RT | GTTTCGCAATTCCTGCAAAT |
| HtrA-R-RT | TGGTGTAGTTGTTCGTTCGG |
